# ER-Mitochondria Contacts Promote Mitochondrial-Derived Compartment Biogenesis

**DOI:** 10.1101/2020.03.13.991133

**Authors:** Alyssa M. English, Benoît Kornmann, Janet M. Shaw, Adam L. Hughes

## Abstract

Mitochondria are dynamic organelles with essential roles in signaling and metabolism. We recently identified a new cellular structure called the mitochondrial-derived compartment (MDC) that is generated from mitochondria in response to amino acid elevation. MDCs protect cells from amino acid toxicity, but how cells form MDCs is unclear. Here, we show that MDCs are micron-sized, lumen-containing organelles that form at sites of contact between the ER and mitochondria. Upon formation, MDCs stably persist at ER-mitochondria contacts for extended periods of time. MDC formation requires the ER-mitochondria encounter structure (ERMES) and GTP hydrolysis by the conserved GTPase Gem1. Unexpectedly, MDC formation is not linked to the role of ERMES/Gem1 in the maintenance of mitochondrial phospholipid homeostasis. Our results identify an important role for ER-mitochondria contacts in the biogenesis of MDCs.

Abbreviations used in this paper: ERMES, ER-mitochondria encounter structure; IMM, inner mitochondrial membrane; MDC, mitochondrial-derived compartment; OMM, outer mitochondrial membrane.

**Summary:** English et al. use super-resolution imaging to show that mitochondrial-derived compartments are lumen-containing organelles that form at sites of contact between the ER and mitochondria. Mitochondrial-derived compartment biogenesis requires a noncanonical function of the ERMES complex and the conserved GTPase Gem1.

## Introduction

Mitochondria are metabolic and signaling organelles that act centrally in ATP production, heme and iron-sulfur cluster synthesis, and the metabolism of lipids, nucleotides, amino acids, and other metabolites (Rutter and Hughes, 2015). Dysfunctional mitochondria are linked to many age-related and metabolic disorders (Nunnari and Suomalainen, 2012; Wallace, 2005). As such, cells are adept at maintaining mitochondrial homeostasis under a variety of metabolic and environmental stress conditions. Well-characterized systems used by cells to maintain mitochondrial homeostasis include mitochondrial fission and fusion machinery (Labbé et al., 2014), mitochondrial-localized proteases (Quirós et al., 2015), mitophagy (Pickles et al., 2018), the ubiquitin-proteasome system (Karbowski and Youle, 2011), mitochondrial-derived vesicles (Sugiura et al., 2014), and the mitochondrial unfolded protein response (Shpilka and Haynes, 2018). These pathways operate in coordination to control mitochondrial health, and failure of these systems is linked to a host of disorders (Nunnari and Suomalainen, 2012; Wallace, 2005).

We recently discovered a new mitochondrial protein remodeling pathway in budding yeast, *S. cerevisiae*, called the mitochondrial-derived compartment (MDC) pathway (Hughes et al., 2016). MDCs are distinct subdomains of mitochondria that are generated in response to defects in lysosomal acidification. These structures selectively incorporate the outer mitochondrial membrane (OMM) import receptor Tom70 and inner mitochondrial membrane (IMM) metabolite carriers of the SLC25 family, while leaving the remainder of the mitochondrial proteome intact. After formation, MDCs are removed from mitochondria by fission and degraded in the yeast lysosome (vacuole) by autophagy. In a separate study currently under review, we found that MDC formation is triggered by a variety of insults that acutely elevate intracellular amino acid pools, including impairment of lysosomal amino acid storage, inhibition of protein translation, exposure to amino acid-derived aldehydes, and inhibition of the mechanistic target of rapamycin pathway (Schuler et al., 2020). We also show that MDC activation is coordinated with metabolite transporter control on other cellular membranes, and that the MDC pathway cooperates with lysosomes and the multi-vesicular body/endosomal sorting complexes required for transport pathway (Henne et al., 2011) to protect cells from elevated amino acid stress. Despite the emerging importance of this pathway in maintaining mitochondrial homeostasis and cell health, it is unclear how cells generate MDCs and selectively sort proteins into them.

Here, we sought to illuminate the mechanism(s) and machinery underlying MDC formation. Using super-resolution imaging, we show that MDCs are dynamic, lumen-containing organelles that generate from mitochondria at sites of contact with the ER. MDC biogenesis at ER-mitochondria contacts requires the ER-mitochondria encounter structure (ERMES) and GTP hydrolysis by the conserved GTPase Gem1. Surprisingly, the role of ERMES/Gem1 in MDC formation appears to be independent of known ERMES/Gem1 functions, as MDCs are not recovered in ERMES/Gem1 mutants rescued by an artificial ER-mitochondria tether or by a suppressor mutation in the endosomal protein Vps13. These results identify a new function for ER-mitochondria contacts in the biogenesis of MDCs, amino acid-responsive organelles.

## Results

### MDCs are dynamic, lumen-containing organelles

We previously identified MDCs as foci associated with mitochondria in cells with dysfunctional lysosomes (Hughes et al., 2016). In a separate manuscript currently under review, we demonstrate that MDCs form in response to high levels of intracellular amino acids (Schuler et al., 2020). We show that rapamycin, a potent inhibitor of the mechanistic target of rapamycin (Heitman et al., 1991), increases intracellular amino acids and activates MDC formation. Rapamycin-induced MDCs are cargo-selective and are identified by their enrichment of Tom70, an OMM import receptor required for the import of mitochondrial carriers (Söllner et al., 1990), and exclusion of other mitochondrial proteins, including a subset of IMM proteins such as Tim50 (Fig. 1 A) (Yamamoto et al., 2002), as well as soluble matrix proteins. To understand how MDCs form, we captured super-resolution images of MDCs generated in response to rapamycin and found that they are large, circular structures that contain distinct lumens commonly reaching ∼1 μm in diameter (Fig. 1 B). Z-series images of an MDC revealed that the diameter of its lumen decreased near the top and bottom until it was no longer visible, suggesting that MDCs are spherical, membrane-bound compartments (Fig. 1 C). Using super-resolution time-lapse imaging to visualize MDC biogenesis, we observed that MDC formation typically began with accumulation of Tom70 at a distinct site on a mitochondrial tubule. Over time, the Tom70-focus grew into a round structure with a clear lumen (Fig. 1 D and Video S1). MDCs remained associated with mitochondrial tubules for extended periods of time and exhibited dynamic behavior, frequently appearing elongated (Fig. 1 D and Video S1).

**Figure 1.**
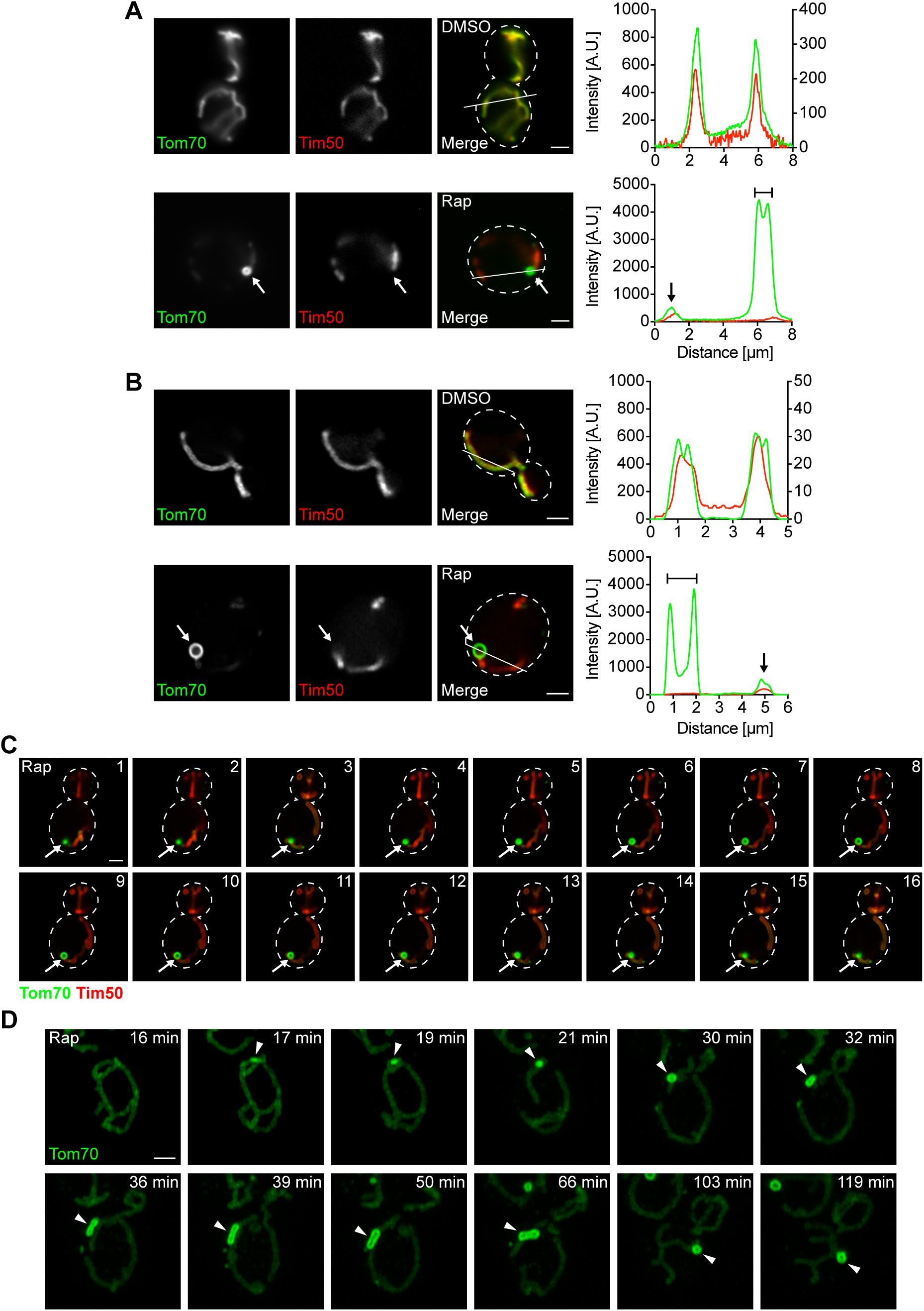
MDCs are dynamic, lumen-containing organelles. (A) Widefield images and line scan analysis of yeast expressing Tom70-GFP and Tim50-mCherry treated with DMSO or rapamycin (Rap). White arrows mark MDC. White line marks fluorescence intensity profile position. Left and right Y axis (top line scan graph) correspond to Tom70-GFP and Tim50-mCherry fluorescence intensity, respectively. Single Y axis (bottom line scan graph) corresponds to Tom70-GFP and Tim50-mCherry fluorescence intensity. Bracket marks MDC. Black arrow marks mitochondrial tubule. Images show single focal plane. Scale bar = 2 µm. (B) Super-resolution images and line scan analysis of yeast expressing Tom70-GFP and Tim50-mCherry treated with DMSO or Rap. White arrows mark MDC. White line marks fluorescence intensity profile position. Left and right Y axis (top line scan graph) correspond to Tom70-GFP and Tim50-mCherry fluorescence intensity, respectively. Single Y axis (bottom line scan graph) corresponds to Tom70-GFP and Tim50-mCherry fluorescence intensity. Bracket marks MDC. Black arrow marks mitochondrial tubule. Images show single focal plane. Scale bar = 2 µm. (C) Z-series of yeast expressing Tom70-GFP and Tim50-mCherry treated with Rap. White arrows mark MDC. Intervals are 0.1 µm. Scale bar = 2 µm. (D) Super-resolution time-lapse images of yeast expressing Tom70-GFP treated with Rap. Arrowheads mark MDC. Images show maximum intensity projection. Scale bar = 2 µm. See also Video S1.

### MDCs form and stably persist at ER-mitochondria contacts

Although the mitochondrial network extends throughout the cell, we typically observed only one MDC per cell upon treatment with rapamycin, suggesting that MDCs are spatially linked to a distinct subcellular location. Events that take place at discrete sites on mitochondria often do so at points of contact with other organelles, such as the ER. We imaged MDCs in yeast expressing the ER marker Sec61-GFP and found that 100% of MDCs associated with ER tubules (Fig. 2 A and Fig. S1 A). MDCs primarily localized adjacent to the peripheral ER (77% of MDCs) and often simultaneously associated with cytoplasmic ER tubules (57% of peripheral ER-associated MDCs, Fig. 2 A and Fig. S1 A).

**Figure 2.**
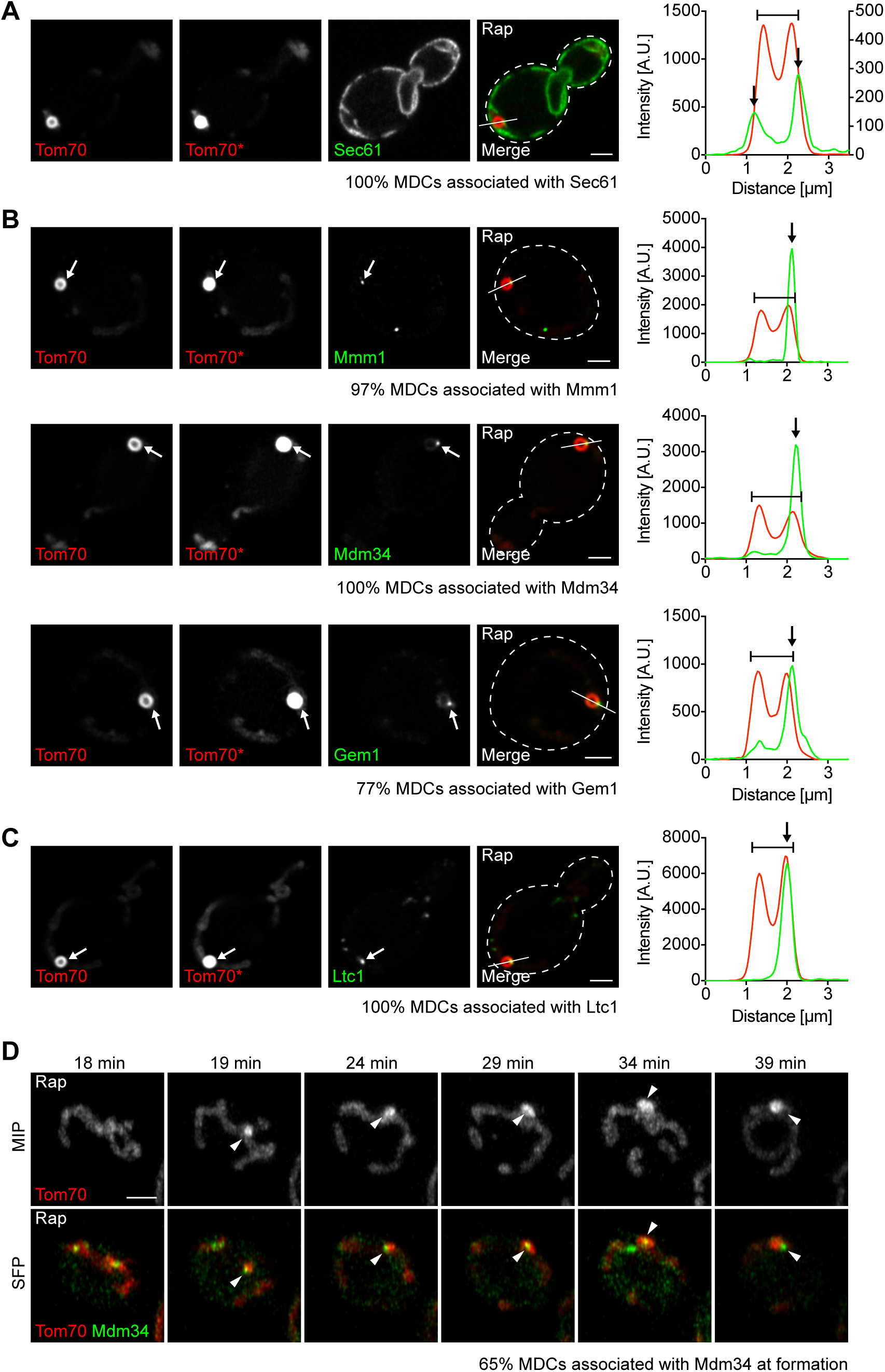
MDCs form and stably persist at ER-mitochondria contacts. (A) Super-resolution images and line scan analysis of yeast expressing Tom70-mCherry and Sec61-GFP treated with rapamycin (Rap). Asterisk (Tom70*) indicates where the fluorescence intensity was increased to visualize mitochondrial tubules. White line marks fluorescence intensity profile position. Left and right Y axis of line scan graph correspond to Tom70-mCherry and Sec61-GFP fluorescence intensity, respectively. Bracket marks MDC. Black arrows mark Sec61 associated with MDC. Images show single focal plane. *n* = 30 MDCs. Scale bar = 2 µm. (B) Super-resolution images and line scan analysis of yeast expressing Tom70-mCherry and Mmm1-GFP, Mdm34-GFP, or Gem1-263GFP treated with Rap. Asterisks (Tom70*) indicate where the fluorescence intensity was increased to visualize mitochondrial tubules. White lines mark fluorescence intensity profile position. Y axis of line scan graph corresponds to Tom70-mCherry and Mmm1-GFP, Mdm34-GFP, or Gem1-263GFP fluorescence intensity. Brackets mark MDCs. White and black arrows mark Mmm1, Mdm34, or Gem1 associated with MDCs. Images show single focal plane. *n* = 30 MDCs. Scale bar = 2 µm. (C) Super-resolution images and line scan analysis of yeast expressing Tom70-mCherry and Ltc1-GFP treated with Rap. Asterisk (Tom70*) indicates where the fluorescence intensity was increased to visualize mitochondrial tubules. White line marks fluorescence intensity profile position. Y axis of line scan graph corresponds to Tom70-mCherry and Ltc1-GFP fluorescence intensity. Bracket marks MDC. White and black arrows mark Ltc1 associated with MDC. Images show single focal plane. *n* = 30 MDCs. Scale bar = 2 µm. (D) Super-resolution time-lapse images of yeast expressing Tom70-mCherry and Mdm34-GFP treated with Rap. Arrowheads mark Mdm34 associated with MDC. Images show maximum intensity projection (MIP) and single focal plane (SFP). *n* = 23 MDCs. Scale bar = 2 µm. See also Fig. S1 and Video S2.

In yeast, mitochondria are tethered to the ER by the ER-mitochondria encounter structure (ERMES), a complex composed of mitochondrial subunits Mdm10, Mdm12, and Mdm34, and the ER subunit Mmm1 (Kornmann et al., 2009). ERMES is regulated by Gem1, a conserved GTPase that colocalizes with the complex (Kornmann et al., 2011; Stroud et al., 2011). Mitochondria and the ER are also connected by the interaction of OMM proteins Tom70 and Tom71 with Ltc1, an ER-resident membrane protein (Elbaz-Alon et al., 2015; Murley et al., 2015). We visualized MDCs in yeast expressing fluorescently-tagged ER-mitochondria contact site proteins, all of which appear as discrete foci simultaneously associated with the ER and mitochondria (Elbaz-Alon et al., 2015; Kornmann et al., 2009, 2011; Murley et al., 2015). MDCs strongly associated with contact site markers Mmm1, Mdm34, Gem1, and Ltc1 (Fig. 2, B and C), indicating that MDCs localize to ER-mitochondria contacts.

We utilized super-resolution time-lapse imaging in cells expressing Mdm34-GFP and Tom70-mCherry to determine whether MDCs formed at ER-mitochondria contacts, or if association with the ER occurred at a later stage of MDC biogenesis. We found that 65% of MDCs associated with Mdm34 at the time of formation (Fig. 2 D and Video S2), and the other 35% associated with Mdm34 within four minutes of initiation. This latter observation suggests that our inability to detect MDC-Mdm34 association in some instances may result from technical limitations. Together, our data indicate that MDC biogenesis is spatially linked to ER-mitochondria contacts and that MDCs remain stably associated with these sites after formation.

### MDC formation requires ERMES and Gem1

We next tested the requirement of ER-mitochondria contact site machinery for MDC formation. We individually deleted genes encoding ERMES subunits Mmm1, Mdm10, Mdm12, and Mdm34, which prevents complex assembly (Kornmann et al., 2009) and produces spherical mitochondria (Berger et al., 1997; Sogo and Yaffe, 1994). We found that these mutants failed to form rapamycin-induced MDCs (Fig. 3, A and B). We also observed inhibition of MDC formation in cells lacking *GEM1* (Fig. 3, A and B). In the absence of Gem1, the ERMES complex remains intact, though *gem1*Δ cells display fewer and larger ERMES foci by fluorescence microscopy (Kornmann et al., 2011). We obtained similar results with other well-characterized MDC inducers, including concanamycin A, which alters lysosomal storage of amino acids by disrupting the activity of the vacuolar H^+^-ATPase (Dröse et al., 1993), and cycloheximide, which increases amino acid pools by inhibiting protein translation (Fig. S2, A and B) (Beugnet et al., 2003). In contrast to ERMES/Gem1, MDC formation was unaltered in *ltc1*Δ cells (Fig. S2 C). MDC biogenesis was also largely unaffected by the loss of other organelle contacts, including vacuole and mitochondria patches (vCLAMPs, *vam6*Δ) (Elbaz-Alon et al., 2014; Hönscher et al., 2014), mitochondria-ER-cortex anchors (MECAs, *num1*Δ) (Lackner et al., 2013), and nucleus-vacuole junctions (NVJs, *nvj1*Δ) (Fig. S2 D) (Pan et al., 2000). These results identify a specific role for ERMES/Gem1 in the biogenesis of MDCs.

**Figure 3.**
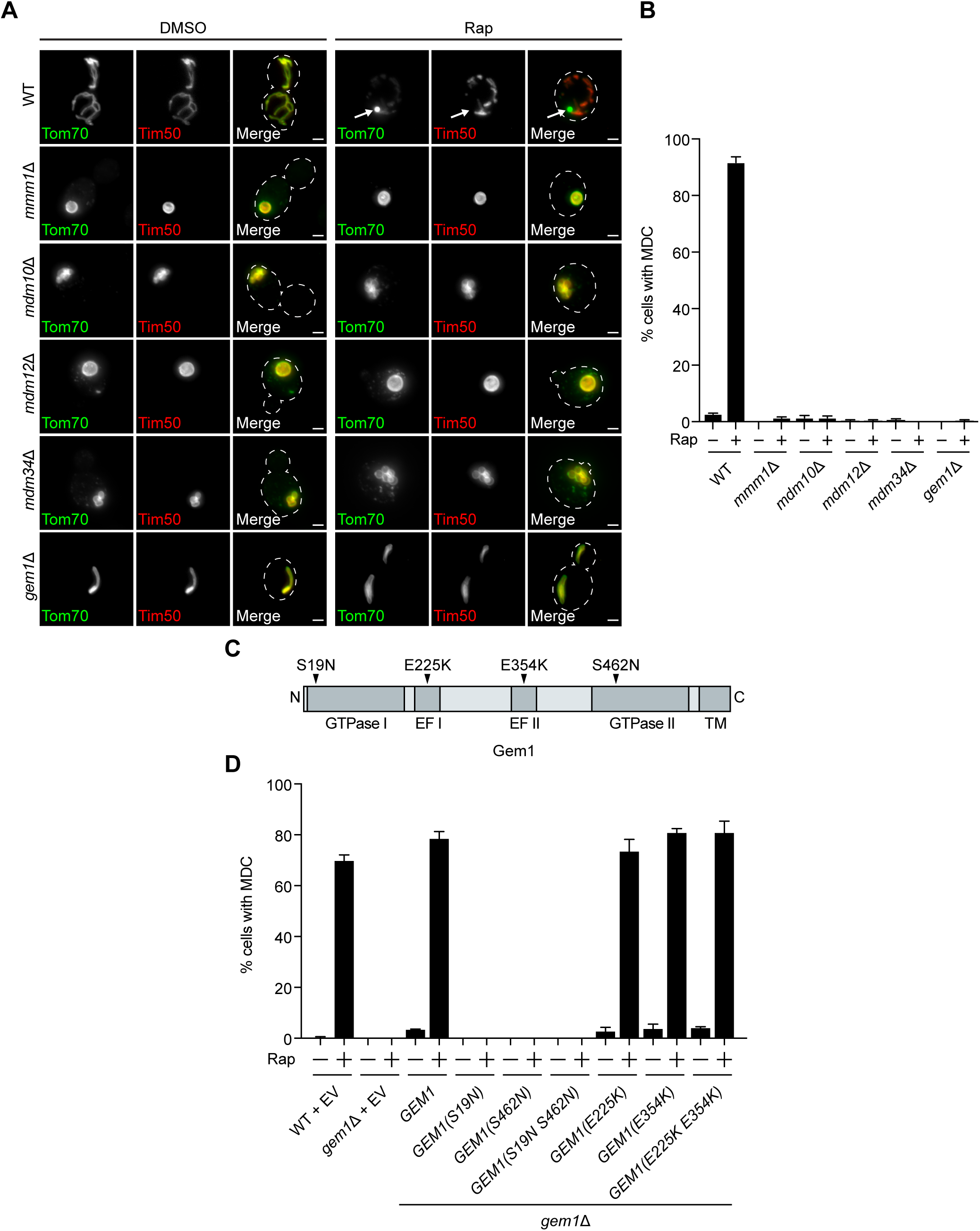
MDC formation requires ERMES and Gem1. (A) Widefield images of wild type (WT) and indicated mutant yeast expressing Tom70-GFP and Tim50-mCherry treated with DMSO or rapamycin (Rap). White arrows mark MDC. Images show maximum intensity projection. Scale bar = 2 µm. (B) Quantification of (A). Error bars show mean ± SE of three replicates, *n* = 100 cells per replicate. (C) Schematic of Gem1 and point mutants used in (D). Transmembrane (TM). (D) Quantification of Rap-induced MDC formation in WT and *gem1*Δ yeast expressing empty vector (EV) or overexpressing *GEM1* or indicated *GEM1* point mutant. Error bars show mean ± SE of three replicates, *n* = 100 cells per replicate. See also Fig. S2.

### GTP hydrolysis by Gem1 is required for MDC formation

Gem1, the yeast ortholog of Miro1/2, is anchored by its C-terminus in the OMM (Fig. 3 C) (Frederick et al., 2004). Gem1 harbors two GTPase domains that hydrolyze GTP and two EF-hands that are capable of binding calcium ions (Fig. 3 C) (Koshiba et al., 2011). To determine if these functional domains were important for MDC formation, we generated well-characterized substitution mutations in the GTPase motifs and EF-hands of Gem1 and overexpressed them in *gem1*Δ cells (Fig. 3 C) (Koshiba et al., 2011). We found that mutations that disrupted Gem1’s ability to bind calcium ions had no effect on MDC formation (Fig. 3 D). However, MDC formation was abolished when Gem1’s ability to hydrolyze GTP via either GTPase domain was compromised (Fig. 3 D). Interestingly, we observed no MDCs when a single GTPase domain was mutated, although single GTPase mutants have reduced, not completely impaired, GTP hydrolysis (Koshiba et al., 2011). Thus, GTP hydrolysis by Gem1 is required for MDC formation.

### The role of ERMES/Gem1 in MDC formation is not linked to mitochondrial phospholipid homeostasis

Next, we aimed to identify the function of ERMES/Gem1 in MDC formation. The central role of the ERMES complex is to facilitate phospholipid transport between mitochondria and the ER (Kornmann et al., 2009). Phospholipid synthesis takes place at mitochondria and the ER, and many phospholipid intermediates are transported between the two organelles. Specifically, phosphatidylserine is generated at the ER and can be transported to the IMM where it is converted to phosphatidylethanolamine by a mitochondria-localized pool of phosphatidylserine decarboxylase Psd1 (Clancey et al., 1993; Friedman et al., 2018; Trotter et al., 1993). Similarly, phosphatidic acid is trafficked from the ER to the IMM by Ups1 (Connerth et al., 2012) and ultimately converted to cardiolipin by Crd1 (Chang et al., 1998; Tuller et al., 1998). Yeast cells lacking ERMES have abnormal phospholipid levels (Kornmann et al., 2009), and ERMES complex subunits Mmm1, Mdm12, and Mdm34 possess lipid-binding domains (AhYoung et al., 2015; Jeong et al., 2016, 2017; Kawano et al., 2018; Kopec et al., 2010). Therefore, we considered it likely that ERMES was required to achieve a proper mitochondrial phospholipid composition necessary for MDC formation. To test this, we assessed MDC formation in various phospholipid synthesis mutants, including *psd1*Δ, *ups1*Δ, and *crd1*Δ. Because yeast contain Psd2, an additional phosphatidylserine decarboxylase that localizes to the Golgi and lysosome (Trotter and Voelker, 1995), we also examined MDC formation in *psd2*Δ and *psd1*Δ *psd2*Δ cells. In contrast to the MDC defects we observed in ERMES/Gem1 mutants, MDC formation was unaffected in phospholipid synthesis mutants (Fig. 4 A), suggesting that the ERMES complex does not function to ensure proper mitochondrial phospholipid composition for MDC biogenesis.

**Figure 4.**
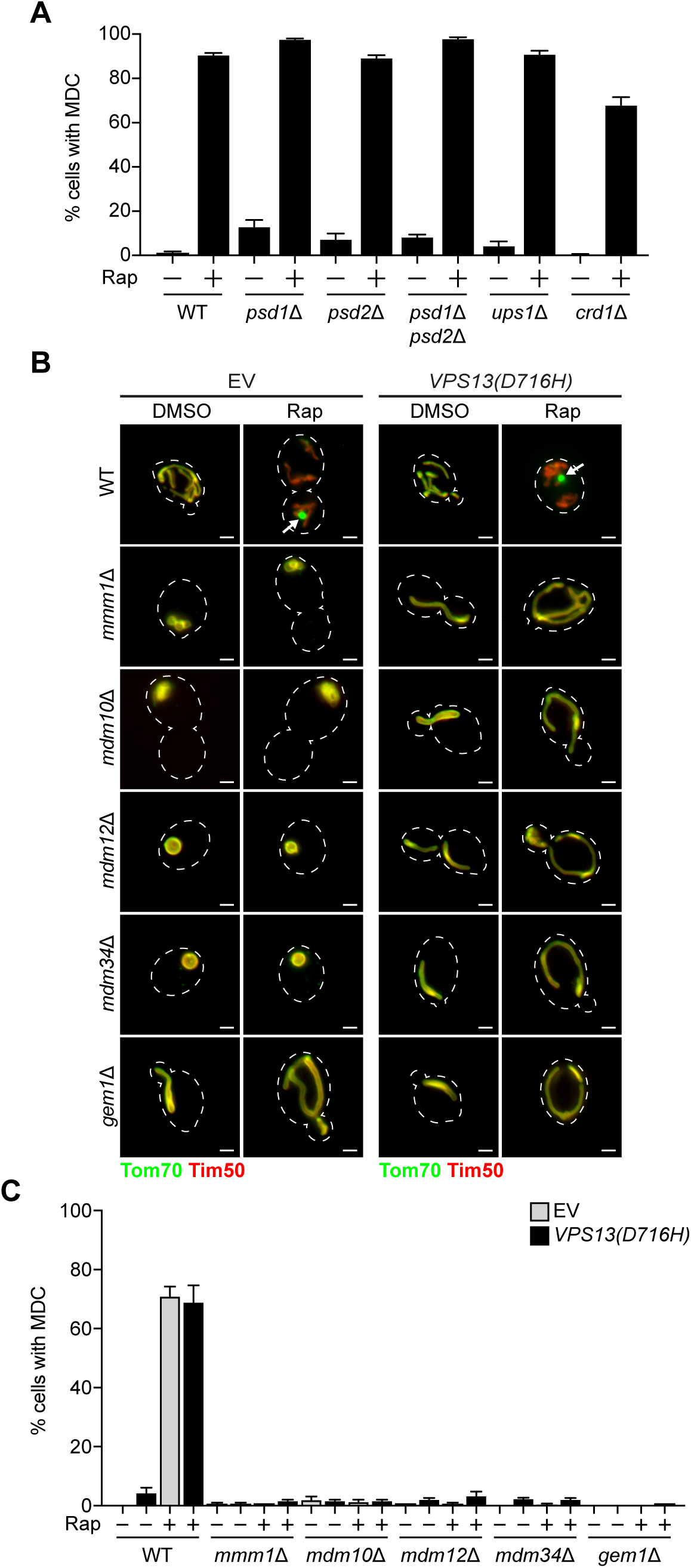
The role of ERMES/Gem1 in MDC formation is not linked to mitochondrial phospholipid homeostasis. (A) Quantification of rapamycin (Rap)-induced MDC formation in wild type (WT) and indicated mutant yeast. Error bars show mean ± SE of three replicates, *n* = 100 cells per replicate. (B) Widefield images of WT and indicated mutant yeast expressing Tom70-GFP and Tim50-mCherry and empty vector (EV) or *VPS13(D716H)* treated with DMSO or Rap. White arrows mark MDCs. Images show maximum intensity projection of merged Tom70-GFP and Tim50-mCherry. Scale bar = 2 µm. (C) Quantification of (B). Error bars show mean ± SE of three replicates, *n* = 100 cells per replicate. See also Fig. S3.

As an alternative approach to test whether the loss of MDCs in ERMES mutants was due to disrupted phospholipid synthesis, we took advantage of a dominant substitution mutation in the endosomal protein Vps13 (Lang et al., 2015). Cells lacking subunits of the ERMES complex characteristically display spherical mitochondria (Berger et al., 1997; Sogo and Yaffe, 1994), loss of mitochondrial DNA (Hobbs et al., 2001), minimal growth on fermentable media, and no growth on nonfermentable media (Berger et al., 1997; Burgess et al., 1994; Youngman et al., 2004). Expression of the *VPS13(D716H)* allele was reported to suppress the characteristic abnormalities exhibited by ERMES mutants, importantly, without restoring complex assembly (Lang et al., 2015). Three lines of evidence suggest that *VPS13(D716H)* suppresses ERMES mutant-associated defects by restoring mitochondrial lipid homeostasis. First, Vps13 localizes to vacuole and mitochondria patches (vCLAMPs) (John Peter et al., 2017), contacts between mitochondria and lysosomes that expand in the absence of ERMES to funnel lipids to mitochondria (Elbaz-Alon et al., 2014; Hönscher et al., 2014). Second, loss of *VPS13* and ERMES is synthetically lethal, suggesting that Vps13 functions redundantly with ERMES (Lang et al., 2015). Third, Vps13 has the capacity to transport lipids between organelles (Kumar et al., 2018). We expressed *VPS13(D716H)* in cells lacking individual ERMES subunits or Gem1 and observed tubular mitochondrial morphology (Fig. 4 B) and wild type-like growth on fermentable and nonfermentable media (Fig. S3 A), consistent with previous findings (Lang et al., 2015; Park et al., 2016). We then assessed MDC formation in these strains and found that, although mitochondrial health was restored, MDCs failed to form in response to rapamycin, concanamycin A, or cycloheximide (Fig. 4, B and C, Fig. S3, B and C).

Finally, we wondered if artificially tethering mitochondria to the ER could restore the ability of ERMES/Gem1 mutants to generate MDCs, though it seemed unlikely as the results from our *VPS13(D716H)* experiments indicated that the presence of the ERMES complex was critical for MDC formation. We utilized ChiMERA, a synthetic ER-mitochondria tether that suppresses the abnormal mitochondrial morphology and growth deficiencies displayed by ERMES mutants (Fig. S3 D) (Kornmann et al., 2009). Like the *VPS13(D716H)* suppression experiments, we were unable to observe MDCs in *mdm12*Δ or *gem1*Δ cells expressing ChiMERA (Fig. S3 E). Altogether, these findings suggest that the function of ERMES/Gem1 in MDC formation is distinct from its canonical role in the maintenance of mitochondrial phospholipid homeostasis.

## Discussion

The data presented here provide a significant step forward in our understanding of the molecular underpinnings of how cells form MDCs, structures with important implications in the maintenance of cellular nutrient homeostasis. Using super-resolution imaging, we now show that MDCs are dynamic, lumen-containing organelles. We also demonstrate that MDC biogenesis is spatially linked to ER-mitochondria contacts and that MDCs mature at these sites over time. Importantly, we identify ERMES/Gem1 as important factors for MDC biogenesis – the first machinery discovered that specifically governs the formation of this unique cellular structure.

These results raise several important questions for future consideration. Chief among them is: what is the role of ERMES/Gem1 and the ER in MDC formation? To date, the best-characterized function of ERMES is to facilitate phospholipid transport between mitochondria and the ER. However, we find that MDCs persist in numerous phospholipid synthesis mutants and that restoration of mitochondrial lipid homeostasis by expression of *VPS13(D716H)* in ERMES/Gem1 mutants does not restore MDC formation. Thus, it seems likely that ERMES/Gem1 provide an uncharacterized function for the biogenesis of MDCs. But what exactly is that function? One distinct possibility is the recruitment of cytoskeletal machinery that may help shape the MDC. The ERMES complex has been implicated in the attachment of mitochondria to the actin cytoskeleton (Boldogh et al., 1998, 2003), and it is likely that the creation of a large cellular structure like the MDC requires a cytoskeletal-based force. Consistent with this idea, Miro1/2, Gem1 orthologs, are well-known to connect mitochondria to microtubules for movement in mammals (Fransson et al., 2006; Glater et al., 2006; Guo et al., 2005). Though Gem1 has not been demonstrated to join mitochondria and the actin cytoskeleton or microtubules, it is possible that it may do so in the context of MDCs.

In addition to the role of ERMES/Gem1 in MDC formation, the exact membrane topology of the MDC remains elusive, as does the sorting mechanism that governs the selection of cargo into the MDC. What has become clear through our work in a separate manuscript currently under review is that MDCs are generated in response to distinct signaling cues – specifically, an increase in intracellular amino acids (Schuler et al., 2020). How metabolic alterations are relayed to the MDC pathway remains a mystery, one that will likely be solved by the identification of additional biogenesis and sorting machinery. As with the discovery of any new cellular system, much remains to be resolved regarding MDCs, and addressing some of these key outstanding questions will be important for elucidating the role of the MDC pathway in the maintenance of cellular amino acid homeostasis.

## Acknowledgements

We thank members of the A.L.H. and J.M.S laboratories for discussion and manuscript comments and L. VanderMeer (Utah) for technical assistance. Research was supported by NIH grants AG043095, GM119694, AG061376, and AG055648 (A.L.H.), NIH T32GM007464 (A.M.E.), NIH GM53466 and GM84970 (J.M.S.), the Howard Hughes Medical Institute (J.M.S.), and the Wellcome Trust 214291/Z/18/Z (B.K.). A.L.H. was further supported by an American Federation for Aging Research Junior Research Grant, United Mitochondrial Disease Foundation Early Career Research Grant, Searle Scholars Award, and Glenn Foundation for Medical Research Award.

## Author contributions

Conceptualization, A.L.H., J.M.S., and A.M.E.; Methodology, A.L.H., J.M.S., B.K., and A.M.E.; Formal Analysis, A.M.E.; Investigation, A.M.E.; Writing - Original Draft, A.M.E.; Writing – Reviewing and Editing, B.K., A.L.H., and J.M.S.; Visualization, A.M.E.; Supervision, A.L.H. and J.M.S.; Funding Acquisition, A.L.H., B.K., J.M.S., and A.M.E.

## Supplemental figure legends

**Figure S1.**
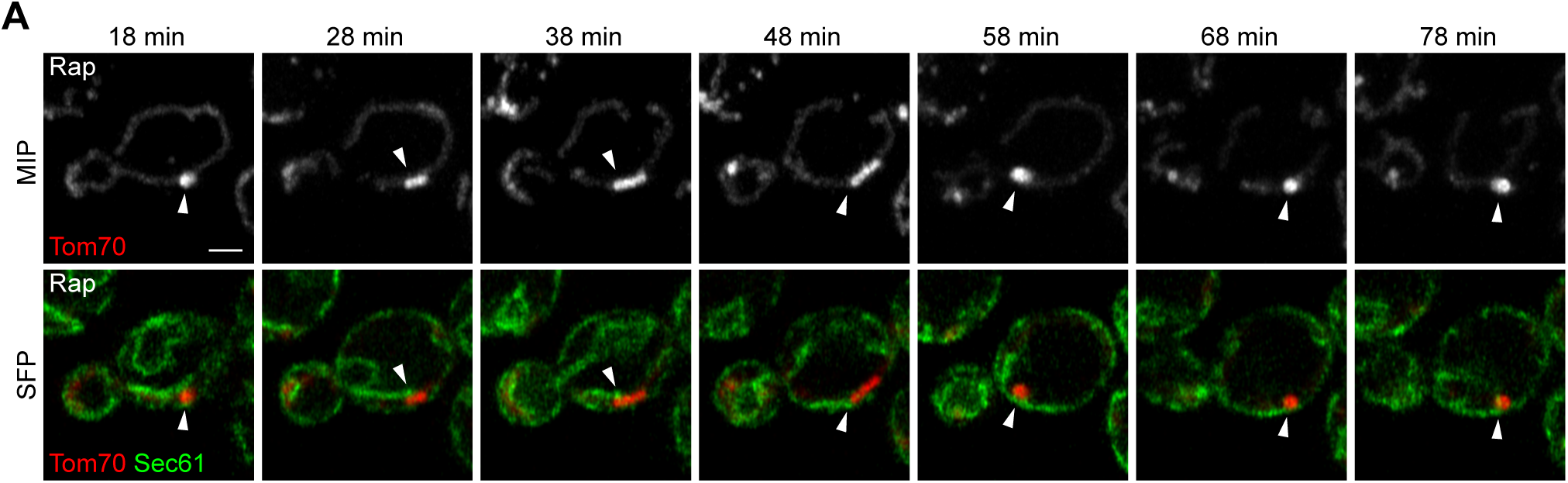
MDCs stably associate with the ER (related to Figure 2). (A) Super-resolution time-lapse images of yeast expressing Tom70-mCherry and Sec61 GFP treated with rapamycin (Rap). Arrowheads mark Sec61 associated with MDC. Images show maximum intensity projection (MIP) and single focal plane (SFP). Scale bar = 2 µm.

**Figure S2.**
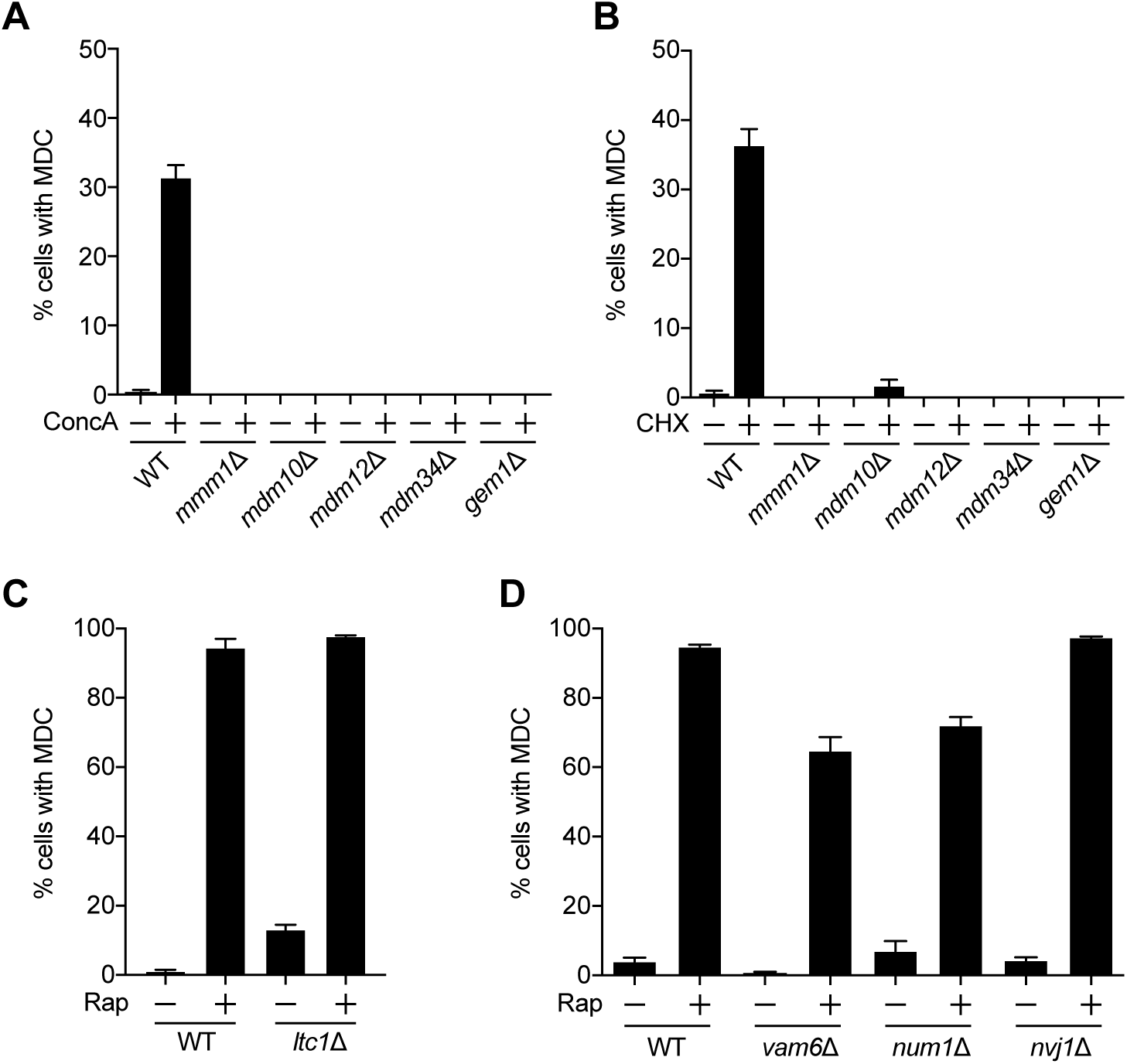
MDC formation requires ERMES and Gem1, but not other organelle contact site proteins (related to Figure 3). (A) Quantification of concanamycin A (ConcA)-induced MDC formation in wild type (WT) and indicated mutant yeast. Error bars show mean ± SE of three replicates, *n* = 100 cells per replicate. (B) Quantification of cycloheximide (CHX)-induced MDC formation in wild type (WT) and indicated mutant yeast. Error bars show mean ± SE of three replicates, *n* = 100 cells per replicate. (C) Quantification of rapamycin (Rap)-induced MDC formation in wild type (WT) and indicated mutant yeast. Error bars show mean ± SE of three replicates, *n* = 100 cells per replicate. (D) Quantification of Rap-induced MDC formation in wild type (WT) and indicated mutant yeast. Error bars show mean ± SE of three replicates, *n* = 100 cells per replicate.

**Figure S3.**
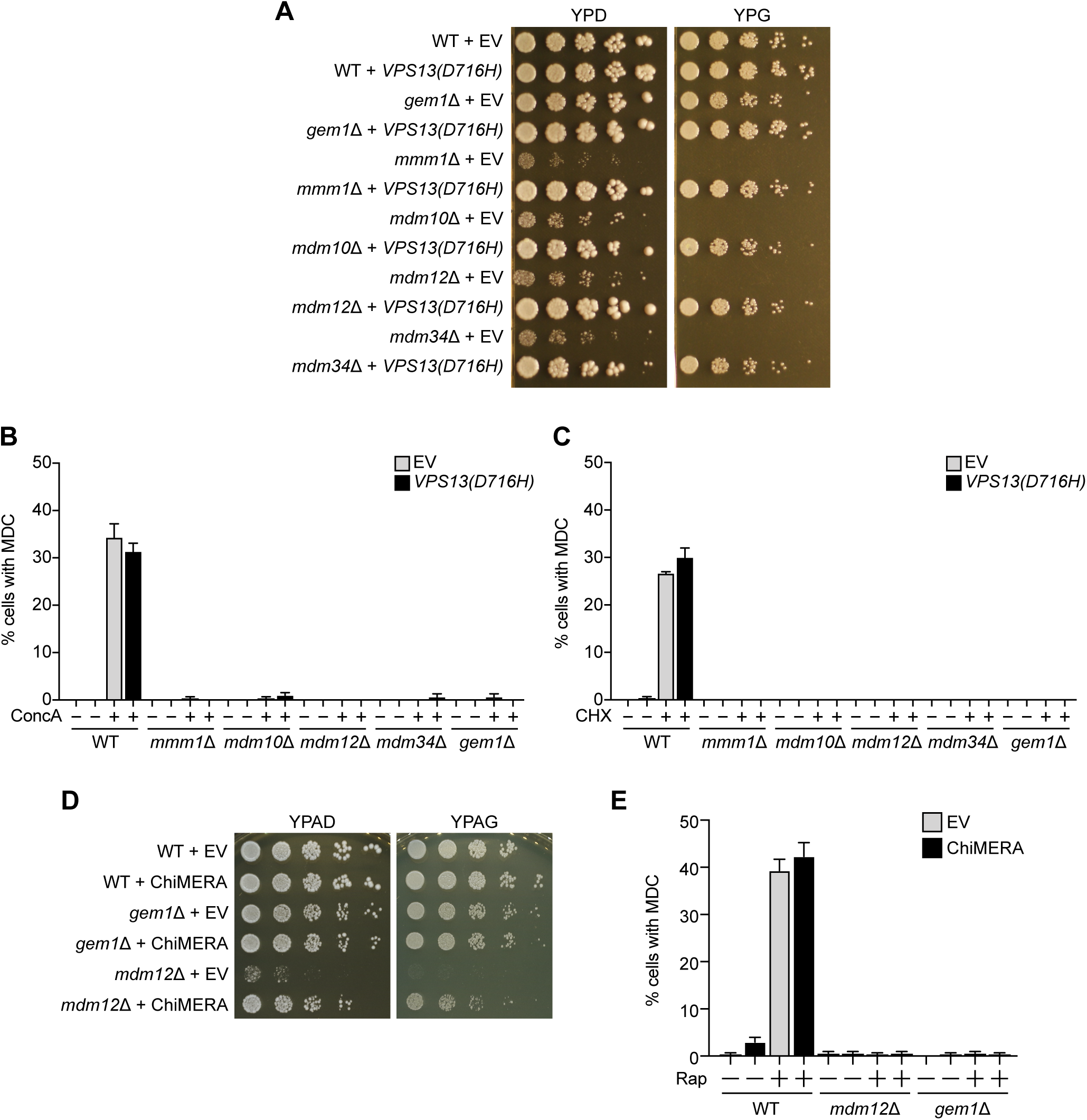
MDC formation in ERMES/Gem1 mutants is not rescued by expression of *VPS13(D716H)* or ChiMERA (related to Figure 4). (A) Serial dilutions of wild type (WT) and indicated mutant yeast expressing empty vector (EV) or *VPS13(D716H)* spotted on media with glucose (YPD) or glycerol (YPG). (B) Quantification of concanamycin A (ConcA)-induced MDC formation in WT and indicated mutant yeast expressing empty vector (EV) or *VPS13(D716H)*. Error bars show mean ± SE of three replicates, *n* = 100 cells per replicate. (C) Quantification of cycloheximide (CHX)-induced MDC formation in WT and indicated mutant yeast expressing EV or *VPS13(D716H)*. Error bars show mean ± SE of three replicates, *n* = 100 cells per replicate. (D) Serial dilutions of WT and indicated mutant yeast expressing EV or ChiMERA spotted on media with glucose (YPAD) or glycerol (YPAG). (E) Quantification of rapamycin (Rap)-induced MDC formation in WT and indicated mutant yeast expressing empty EV or ChiMERA. Error bars show mean ± SE of three replicates, *n* = 100 cells per replicate.

## Supplemental item legends

**Video S1. MDCs form from a single site and are dynamic (related to Fig. 1).**

Maximum intensity projection video of yeast expressing Tom70-GFP treated with rapamycin. Images were taken every minute and are shown at 2 frames per second.

**Video S2. MDCs form at ER-Mitochondria contacts (related to Fig. 2).**

Maximum intensity projection video of yeast expressing Tom70-mCherry and Mdm34-GFP treated with rapamycin. Images were taken every minute and are shown at 2 frames per second.

**Table S1. Yeast strains used in this study.**

**Table S2. Oligonucleotides used in this study.**

## Materials and methods

### Yeast strains and culture

All yeast strains are derivatives of *Saccharomyces cerevisiae* S288C (BY) (Brachmann et al., 1998) and are listed in Table S1. Strains expressing fluorescently-tagged *TOM70*, *TIM50*, *SEC61*, *MMM1*, *MDM34*, and *LTC1* were created by one step PCR-mediated C-terminal endogenous epitope tagging using standard techniques and oligo pairs listed in Table S2. Plasmid templates for fluorescent epitope tagging were from the pKT series of vectors (Sheff and Thorn, 2004). Correct integrations were confirmed by a combination of colony PCR across the chromosomal insertion site and correctly localized expression of the fluorophore by microscopy. Because N-terminal tagging of Gem1 yields only partially functional proteins, yEGFP-tagged *GEM1* was created by PCR-mediated internal endogenous yEGFP tagging after amino acid 262 using the Gauss toolbox (Gauss et al., 2005). Functionality of the fusion protein was assessed by quantifying the shape of segmented mitochondria using the circularity metric (4π*area/perimeter^2^, where 1 is a perfect circle). While wild type mitochondria have a circularity of 0.2 ± 0.02 (mean ± SEM), those of yeast expressing N-terminally tagged Gem1 rose to 0.37 ± 0.02. Mitochondria of yeast expressing internally-tagged Gem1 had a circularity of 0.23 ± 0.02, indicating near wild type shape, and thus functionality of the fusion protein. Functionality of the fusion protein was also assessed by the ability of the cells to form MDCs. Deletion strains were created by one-step PCR-mediated gene replacement using the previously described pRS series of vectors (Brachmann et al., 1998; Sikorski and Hieter, 1989) and oligo pairs listed in Table S2. Correct gene deletions were confirmed by colony PCR across the chromosomal insertion site. For strains bearing deletion of *MMM1*, *MDM10*, *MDM12*, or *MDM34*, one copy of the gene was deleted to create a heterozygous diploid, which was subsequently sporulated to obtain haploid mutants. For Figure 3D, strains expressing empty vector, *GEM1*, *GEM1(S19N*), *GEM1(S462N)*, *GEM1(S19N S462N)*, *GEM1(E225K)*, *GEM1(E354K)*, or *GEM1(E225K E354K)* from a GPD promoter integrated into an empty region of chromosome 1 (199456-199457) were constructed by transformation of parental yeast strains with NotI-digested pAG306GPD-empty chr 1, pAG306GPD-*GEM1* chr 1, pAG306GPD-*GEM1(S19N)* chr 1, pAG306GPD-*GEM1(S462N)* chr 1, pAG306GPD-*GEM1(S19N S462N)* chr 1, pAG306GPD-*GEM1(E225K)* chr 1, pAG306GPD-*GEM1(E354K)* chr 1, or pAG306GPD-*GEM1(E225K E354K)* chr 1, respectively, as previously described (Hughes and Gottschling, 2012).

Yeast cells were grown exponentially for 15-16 hours at 30°C to a final density of 2-7×10^6^ cells/mL before the start of all experiments. This period of overnight log-phase growth was carried out to ensure vacuolar and mitochondrial uniformity across the cell population and is essential for consistent MDC activation. Unless otherwise indicated, cells were cultured in YPAD medium (1% yeast extract, 2% peptone, 0.005% adenine, 2% glucose). Cells expressing pRS315 (Sikorski and Hieter, 1989), p*VPS13(D716H)* (Lang et al., 2015), pRS415 (Brachmann et al., 1998), or p415GPD-ChiMERA (Kornmann et al., 2009) were cultured in SD-leucine medium (0.67% yeast nitrogen base without amino acids, 2% glucose, supplemented nutrients 0.072 g/L each adenine, alanine, arginine, asparagine, aspartic acid, cysteine, glutamic acid, glutamine, glycine, histidine, myo-inositol, isoleucine, lysine, methionine, phenylalanine, proline, serine, threonine, tryptophan, tyrosine, uracil, valine, 0.007 g/L para-aminobenzoic acid) for selection of the plasmids, and, for MDC assays, were switched to YPAD at the time of drug treatment. Unless otherwise indicated, rapamycin (LC Laboritories, R-5000), concanamycin A (Santa Cruz Biotechnology, sc-202111), and cycloheximide (Sigma-Aldrich, C1988) were added to cultures at final concentrations of 200 nM, 500 nM, and 50 μg/mL, respectively.

### Plasmids

To create pAG306GPD-*GEM1* chr 1, we used the donor Gateway plasmid pDONR221-*GEM1* to insert *GEM1* into pAG306GPD-ccdB chr 1 (Hughes and Gottschling, 2012) using LR clonase (Thermo Fisher, 11791020) according to the manufacturer’s instructions. To create pAG306GPD-*GEM1(S19N)* chr 1, pAG306GPD-*GEM1(S462N)* chr 1, pAG306GPD-*GEM1(S19N S462N)* chr 1, pAG306GPD-*GEM1(E225K)* chr 1, pAG306GPD-*GEM1(E354K)* chr 1, and pAG306GPD-*GEM1(E225K E354K)* chr 1, we used the Q5 Site-Directed Mutagenesis Kit (New England Biolabs, E0554S) and pDONR221-*GEM1* to make pDONR221-*GEM1(S19N)*, pDONR221-*GEM1(S462N)*, pDONR221-*GEM1(S19N S462N)*, pDONR221-*GEM1(E225K)*, pDONR221-*GEM1(E354K)*, and pDONR221-*GEM1(E225K E354K)*, respectively. We then used pDONR221-*GEM1(S19N)*, pDONR221-*GEM1(S462N)*, pDONR221-*GEM1(S19N S462N)*, pDONR221-*GEM1(E225K)*, pDONR221-*GEM1(E354K)*, and pDONR221-*GEM1(E225K E354K)* to insert *GEM1(S19N)*, *GEM1(S462N)*, *GEM1(S19N S462N)*, *GEM1(E225K)*, *GEM1(E354K)*, and *GEM1(E225K E354K)*, respectively, into pAG306GPD-ccdB chr 1 using LR clonase according to the manufacturer’s instructions.

### MDC assays

For MDC assays, overnight log-phase cell cultures were grown in the presence of dimethyl sulfoxide (DMSO) (Sigma-Aldrich, D2650) or the indicated drug for two hours. After incubation, cells were harvested by centrifugation and optical z-sections of live yeast cells were acquired with a ZEISS Axio Imager M2 or, for super-resolution images, a ZEISS LSM800 with Airyscan or ZEISS LSM880 with Airyscan. The percent cells with MDCs was quantified in each experiment at the two-hour time point. All quantifications show the mean ± standard error from three biological replicates with *n* = 100 cells per experiment. MDCs were identified as Tom70-positive, Tim50-negative spherical structures that were enriched for Tom70 versus the mitochondrial tubule.

### Microscopy

Optical z-sections of live yeast cells were acquired with a ZEISS Axio Imager M2 equipped with a ZEISS Axiocam 506 monochromatic camera, 100× oil-immersion objective (plan apochromat, NA 1.4), a ZEISS LSM800 equipped with an Airyscan detector, 63× oil-immersion objective (plan apochromat, NA 1.4) or a ZEISS LSM880 equipped with an Airyscan detector, 63× oil-immersion objective (plan apochromat, NA 1.4). Widefield images were acquired with ZEN (Carl Zeiss) and processed with Fiji (Schindelin et al., 2012). Super-resolution images were acquired with ZEN (Carl Zeiss) and processed using the automated Airyscan processing algorithm in ZEN (Carl Zeiss) and Fiji. Individual channels of all images were minimally adjusted in Fiji to match the fluorescence intensities between channels for better visualization. Line scan analysis was performed on non-adjusted, single z-sections in Fiji.

### Time-lapse imaging

For Figures 1D, 2D, S1A, and Videos S1 and S2, overnight log-phase cultures were treated with 1 μM rapamycin for 15 minutes. Cells were harvested by centrifugation, resuspended in SD medium (0.67% yeast nitrogen base without amino acids, 2% glucose, supplemented nutrients 0.074 g/L each adenine, alanine, arginine, asparagine, aspartic acid, cysteine, glutamic acid, glutamine, glycine, histidine, myo-inositol, isoleucine, lysine, methionine, phenylalanine, proline, serine, threonine, tryptophan, tyrosine, uracil, valine, 0.369 g/L leucine, 0.007 g/L para-aminobenzoic acid) and 5 μM rapamycin, and pipetted into flow chamber slides as previously described (Fees et al., 2017). Briefly, flow chambers were made using standard microscope slides and coverslips. Strips of Parafilm were used to seal a coverslip to a slide and created the walls of the chamber. Flow chambers were coated with concanavalin A (Sigma-Aldrich, L7647) prior to loading cells. Melted Vaseline was used to seal the chamber. Optical z-sections of live yeast cells were acquired with a ZEISS Airyscan LSM880 equipped with an environmental chamber set to 30°C. Super-resolution time-lapse images were acquired with ZEN (Carl Zeiss) and processed using the automated Airyscan processing algorithm in ZEN (Carl Zeiss) and Fiji. Individual channels of all images were minimally adjusted in Fiji to match the fluorescence intensities between channels for better visualization.

### Serial-dilution growth assays

Five-fold serial dilutions of exponentially growing yeast cells were in water and 3 μl of each dilution was spotted onto YPD (1% yeast extract, 2% peptone, 2% glucose) and YPG (1% yeast extract, 2% peptone, 3% glycerol) agar, or YPAD (1% yeast extract, 2% peptone, 0.005% adenine, 2% glucose) and YPAG (1% yeast extract, 2% peptone, 0.005% adenine, 3% glycerol) agar. Total cells plated in each dilution spot were 5000, 1000, 20, 40, and 8.

### Quantification and statistical analysis

The number of replicates, what *n* represents, and dispersion and precision measures are indicated in the figure legends. In general, quantifications show the mean ± standard error from three biological replicates with *n* = 100 cells per experiment. In experiments with data depicted from a single biological replicate, the experiment was repeated with the same results.

## Supplemental material

Fig. S1 shows time-lapse images of an MDC stably associating with the ER over time. Fig. S2 demonstrates that ERMES/Gem1 are required for concanamycin A or cycloheximide-induced MDC formation, and other organelle contacts including vacuole and mitochondria patches (vCLAMPs), mitochondria-ER-cortex anchors (MECAs), and nucleus-vacuole junctions (NVJs) are not required for rapamycin-induced MDC formation. Fig. S3 demonstrates that ERMES/Gem1 mutants expressing *VPS13(D716H)* or ChiMERA do not form MDCs in response to concanamycin A, cycloheximide, or rapamycin. Video S1 shows the process of MDC biogenesis. Video S2 shows an MDC forming at and associating with an ER-mitochondria contact site. Table S1 lists the yeast strains used in this study. Table S2 lists the oligonucleotides used in this study.

