## Supplemental Table 1 for "ER-Mitochondria Contacts Promote Mitochondrial-Derived Compartment Biogenesis"

**Table S1. Yeast strains used in this study.**

| Strain | Genotype |
|---|---|
| | BY4741 MATa his3Δ1 leu2Δ0 ura3Δ0 met15Δ0 |
| | BY4742 MATα his3Δ1 leu2Δ0 ura3Δ0 lys2Δ0 |
| | BY4743 MATa/MATα his3Δ1/his3Δ1 leu2Δ0/leu2Δ0 ura3Δ0/ura3Δ0 met15Δ0/+ lys2Δ0/+ |
| AHY1480 | BY4743 TOM70-yEGFP:KanMX/+ TIM50-mCherry:KanMX/+ |
| AHY7053 | BY4743 TOM70-yEGFP:SpHIS5MX/TOM70-yEGFP:SpHIS5MX TIM50-mCherry:KanMX/TIM50-mCherry:KanMX |
| AHY5082 | BY4743 TOM70-yEGFP:KanMX/TOM70-yEGFP:KanMX |
| AHY7358 | BY4743 TOM70-mCherry:KanMX/TOM70-mCherry:KanMX SEC61-yEGFP:SpHIS5MX/SEC61-yEGFP:SpHIS5MX |
| AHY3694 | BY4743 TOM70-mCherry:KanMX/TOM70-mCherry:KanMX MMM1-yEGFP:SpHIS5MX/MMM1-yEGFP:SpHIS5MX |
| AHY3696 | BY4743 TOM70-mCherry:KanMX/TOM70-mCherry:KanMX MDM34-yEGFP:SpHIS5MX/MDM34-yEGFP:SpHIS5MX |
| ByK302 | BY4741 MDM34-mCherry:SpHIS5MX GEM1-263yEGFP pVTU100-mtBFP |
| AHY7499 | BY4743 TOM70-mCherry:KanMX/TOM70-mCherry:KanMX GEM1-263yEGFP/GEM1-263yEGFP |
| AHY3698 | BY4743 TOM70-mCherry:KanMX/TOM70-mCherry:KanMX LTC1-yEGFP:spHIS5MX/LTC1-yEGFP:spHIS5MX |
| AHY4057 | BY4743 TOM70-yEGFP:KanMX/+ TIM50-mCherry:KanMX/+ gem1Δ::HygMX/gem1Δ::HygMX |
| AHY4061 | TOM70-yEGFP:KanMX/+ TIM50-mCherry:KanMX/+ mmm1Δ::HygMX/mmm1Δ::HygMX his3Δ1/his3Δ1 leu2Δ0/leu2Δ0 ura3Δ0/ura3Δ0 met15Δ0/met15Δ0 |
| AHY4082 | BY4743 TOM70-yEGFP:KanMX/+ TIM50-mCherry:KanMX/+ mdm10Δ::HygMX/mdm10Δ::HygMX |

| | |
|---|---|
| AHY4221 | TOM70-yEGFP:KanMX/+ TIM50-mCherry:KanMX/+ mdm12Δ::HygMX/mdm12Δ::HygMX his3Δ1/his3Δ1 leu2Δ0/leu2Δ0 ura3Δ0/ura3Δ0 met15Δ0/+ lys2Δ0/lys2Δ0 |
| AHY4167 | TOM70-yEGFP:KanMX/+ TIM50-mCherry:KanMX/+ mdm34Δ::HygMX/mdm34Δ::HygMX his3Δ1/his3Δ1 leu2Δ0/leu2Δ0 ura3Δ0/ura3Δ0 met15Δ0/+ lys2Δ0/lys2Δ0 |
| AHY4046 | BY4743 TOM70-yEGFP:KanMX/+ TIM50-mCherry:KanMX/+ ltc1Δ::URA3/ltc1Δ::URA3 |
| AHY1938 | BY4743 TOM70-yEGFP:KanMX/+ TIM50-mCherry:KanMX/+ vam6Δ::URA3/vam6Δ::URA3 |
| AHY9563 | BY4743 TOM70-yEGFP:KanMX/+ TIM50-mCherry:KanMX/+ num1Δ::URA3/num1Δ::URA3 |
| AHY4050 | BY4743 TOM70-yEGFP:KanMX/+ TIM50-mCherry:KanMX/+ nvj1Δ::URA3/nvj1Δ::URA3 |
| AHY8245 | BY4743 TOM70-yEGFP:KanMX/+ TIM50-mCherry:KanMX/+ chr 1(199456-199457)::$P_{GPD1}$-$Term_{CYC1}$-URA3/+ |
| AHY8247 | BY4743 TOM70-yEGFP:KanMX/+ TIM50-mCherry:KanMX/+ gem1Δ::HygMX/gem1Δ::HygMX chr 1(199456-199457)::$P_{GPD1}$-$Term_{CYC1}$-URA3/+ |
| AHY8136 | BY4743 TOM70-yEGFP:KanMX/+ TIM50-mCherry:KanMX/+ gem1Δ::HygMX/gem1Δ::HygMX chr 1(199456-199457)::$P_{GPD1}$-GEM1-$Term_{CYC1}$-URA3/+ |
| AHY8377 | BY4743 TOM70-yEGFP:KanMX/+ TIM50-mCherry:KanMX/+ gem1Δ::HygMX/gem1Δ::HygMX chr 1(199456-199457)::$P_{GPD1}$-GEM1(S19N)-$Term_{CYC1}$-URA3/+ |
| AHY8383 | BY4743 TOM70-yEGFP:KanMX/+ TIM50-mCherry:KanMX/+ gem1Δ::HygMX/gem1Δ::HygMX chr 1(199456-199457)::$P_{GPD1}$-GEM1(S462N)-$Term_{CYC1}$-URA3/+ |
| AHY8385 | BY4743 TOM70-yEGFP:KanMX/+ TIM50-mCherry:KanMX/+ gem1Δ::HygMX/gem1Δ::HygMX chr 1(199456-199457)::$P_{GPD1}$-GEM1(S19N S462N)-$Term_{CYC1}$-URA3/+ |

| | |
|---|---|
| AHY8379 | BY4743 TOM70-yEGFP:KanMX/+ TIM50-mCherry:KanMX/+ gem1Δ::HygMX/gem1Δ::HygMX chr 1(199456-199457)::$P_{GPD1}$-GEM1(E225K)-Term$_{CYC1}$-URA3/+ |
| AHY8381 | BY4743 TOM70-yEGFP:KanMX/+ TIM50-mCherry:KanMX/+ gem1Δ::HygMX/gem1Δ::HygMX chr 1(199456-199457)::$P_{GPD1}$-GEM1(E354K)-Term$_{CYC1}$-URA3/+ |
| AHY8387 | BY4743 TOM70-yEGFP:KanMX/+ TIM50-mCherry:KanMX/+ gem1Δ::HygMX/gem1Δ::HygMX chr 1(199456-199457)::$P_{GPD1}$-GEM1(E225K E354K)-Term$_{CYC1}$-URA3/+ |
| AHY2423 | BY4743 TOM70-yEGFP:KanMX/+ TIM50-mCherry:KanMX/+ psd1Δ::URA3/psd1Δ::URA3 |
| AHY4671 | BY4743 TOM70-yEGFP:KanMX/+ TIM50-mCherry:KanMX/+ psd2Δ::LEU2/psd2Δ::LEU2 |
| AHY4673 | BY4743 TOM70-yEGFP:KanMX/+ TIM50-mCherry:KanMX/+ psd1Δ::URA3/psd1Δ::URA3 psd2Δ::LEU2/psd2Δ::LEU2 |
| AHY2425 | BY4743 TOM70-yEGFP:KanMX/+ TIM50-mCherry:KanMX/+ ups1Δ::URA3/ups1Δ::URA3 |
| AHY2421 | BY4743 TOM70-yEGFP:KanMX/+ TIM50-mCherry:KanMX/+ crd1Δ::URA3/crd1Δ::URA3 |
| AHY4482 | BY4743 TOM70-yEGFP:KanMX/+ TIM50-mCherry:KanMX/+ pRS315 |
| AHY4484 | BY4743 TOM70-yEGFP:KanMX/+ TIM50-mCherry:KanMX/+ pVPS13(D716H) |
| AHY4490 | BY4743 TOM70-yEGFP:KanMX/+ TIM50-mCherry:KanMX/+ gem1Δ::HygMX/gem1Δ::HygMX pRS315 |
| AHY4492 | BY4743 TOM70-yEGFP:KanMX/+ TIM50-mCherry:KanMX/+ gem1Δ::HygMX/gem1Δ::HygMX pVPS13(D716H) |
| AHY4558 | TOM70-yEGFP:KanMX/+ TIM50-mCherry:KanMX/+ mmm1Δ::HygMX/mmm1Δ::HygMX his3Δ1/his3Δ1 leu2Δ0/leu2Δ0 ura3Δ0/ura3Δ0 met15Δ0/met15Δ0 pRS315 |
| AHY6782 | BY4743 TOM70-yEGFP:KanMX/+ TIM50-mCherry:KanMX/+ mmm1Δ::HygMX/mmm1Δ::HygMX pVPS13(D716H) |

| | |
|---|---|
| AHY4555 | BY4743 TOM70-yEGFP:KanMX/+ TIM50-mCherry:KanMX/+ mdm10Δ::HygMX/mdm10Δ::HygMX pRS315 |
| AHY4503 | TOM70-yEGFP:KanMX/+ TIM50-mCherry:KanMX/+ mdm10Δ::HygMX/mdm10Δ::HygMX his3Δ1/his3Δ1 leu2Δ0/leu2Δ0 ura3Δ0/ura3Δ0 pVPS13(D716H) |
| AHY4572 | TOM70-yEGFP:KanMX/+ TIM50-mCherry:KanMX/+ mdm12Δ::HygMX/mdm12Δ::HygMX his3Δ1/his3Δ1 leu2Δ0/leu2Δ0 ura3Δ0/ura3Δ0 met15Δ0/+ lys2Δ0/lys2Δ0 pRS315 |
| AHY4549 | TOM70-yEGFP:KanMX/+ TIM50-mCherry:KanMX/+ mdm12Δ::HygMX/mdm12Δ::HygMX his3Δ1/his3Δ1 leu2Δ0/leu2Δ0 ura3Δ0/ura3Δ0 met15Δ0/met15Δ0 lys2Δ0/lys2Δ0 pVPS13(D716H) |
| AHY4612 | TOM70-yEGFP:KanMX/+ TIM50-mCherry:KanMX/+ mdm34Δ::HygMX/mdm34Δ::HygMX his3Δ1/his3Δ1 leu2Δ0/leu2Δ0 ura3Δ0/ura3Δ0 met15Δ0/+ lys2Δ0/lys2Δ0 pRS315 |
| AHY4528 | TOM70-yEGFP:KanMX/+ TIM50-mCherry:KanMX/+ mdm34Δ::HygMX/mdm34Δ::HygMX his3Δ1/his3Δ1 leu2Δ0/leu2Δ0 ura3Δ0/ura3Δ0 met15Δ0/met15Δ0 lys2Δ0/lys2Δ0 pVPS13(D716H) |
| AHY4867 | BY4743 TOM70-mCherry:KanMX/TOM70-mCherry:KanMX pRS415 |
| AHY4875 | BY4743 TOM70-mCherry:KanMX/TOM70-mCherry:KanMX p415GPD-ChiMERA |
| AHY4869 | BY4743 TOM70-mCherry:KanMX/TOM70-mCherry:KanMX gem1Δ::HygMX/gem1Δ::HygMX pRS415 |
| AHY4877 | BY4743 TOM70-mCherry:KanMX/TOM70-mCherry:KanMX gem1Δ::HygMX/gem1Δ::HygMX p415GPD-ChiMERA |
| AHY6843 | BY4743 TOM70-mCherry:KanMX/TOM70-mCherry:KanMX mdm12Δ::HygMX/mdm12Δ::HygMX pRS415 |
| AHY6730 | TOM70-mCherry:KanMX/TOM70-mCherry:KanMX mdm12Δ::HygMX/mdm12Δ::HygMX his3Δ1/his3Δ1 leu2Δ0/leu2Δ0 ura3Δ0/ura3Δ0 lys2Δ0/lys2Δ0 p415GPD-ChiMERA |
