## Supplemental Table 2 for "ER-Mitochondria Contacts Promote Mitochondrial-Derived Compartment Biogenesis"

**Table S2. Oligonucleotides used in this study.**

| Name | Sequence |
|---|---|
| Tom70 Tag F | TCAAGAAACTTTAGCTAAATTACGCGAACAGGGTTTAATGGGTGACGGTGCTGGTTTA |
| Tom70 Tag R | TTTGTCTTCTCCTAAAAGTTTTTAAGTTTATGTTTACTGTTCGATGAATTCGAGCTCG |
| Tim50 Tag F | TGAAGAGGAAAAGAAAAAGAAGAAGATTGCTGAATCCAAAGGTGACGGTGCTGGTTTA |
| Tim50 Tag R | ATAGATACGTAGATACATGAGAAGAGGGTTTACATGAAAATCGATGAATTCGAGCTCG |
| Sec61 Tag F | GTTTACTAAGAACCTCGTTCCAGGATTTTCTGATTTGATGggtgacggtgctggttta |
| Sec61 Tag R | GCGATTTTTTTTTTCTTTGGATATTATTTTCATTTTATATtcgatgaattcgagctcg |
| Mmm1 Tag F | ACGTAGTAAAAATACGAGAGAAGAAAAGCCTACAGAGTTAggtgacggtgctggttta |
| Mmm1 Tag R | GATAGGAAAAAGATAGAACAAAAAATTTGTACATAAATATtcgatgaattcgagctcg |
| Mdm34 Tag F | CTGGAAATGGGGCATGGAGGATAGCCCCCCACCATATCATggtgacggtgctggttta |
| Mdm34 Tag R | ATGTATTTGTGTAGTTATGTACTTAGATATGTAACTTAATtcgatgaattcgagctcg |
| Ltc1 Tag F | CAA AGT ACA ATG GTC AAG ACC AAG AAG GGG AGA CGT GTT GGG TGA CGG TGC TGG TTT A |
| Ltc1 Tag R | ACA TAC ATA ACC TTA CAT ACA ATA TTA CAT GAT TAC TCT GTC GAT GAA TTC GAG CTC G |
| Gem1 KO F | AAATAGCGGACTTCTAAATACTAATGTGTTGAACAACACAGATTGTACTGAGAGTGCACC |
| Gem1 KO R | GAAATGCAACACTTCCCTAATATAGAAATTTGGGCATTAACTGTGCGGTATTTCACACCG |

| | |
|---|---|
| Gem1 KO Check | GAGCAGCCACATCAACAGGGCAATGAATTA |
| Mmm1 KO F | TTGAGAGAGTCAATATAATACCTGTAGCCTTTTTCTGAAAGATTGTACTGAGAGTGCACC |
| Mmm1 KO R | GATAGGAAAAAGATAGAACAAAAAATTTGTACATAAATATCTGTGCGGTATTTCACACCG |
| Mmm1 KO Check | TTGATTGTGCACTCGTAAGTGACTTGACTG |
| Mdm10 KO F | TACGTTAGGAAAAAGACACGAACAGAGAAGACCGATCTTGGATTGTACTGAGAGTGCACC |
| Mdm10 KO R | ACCTGTATATTAAAACCTTTATTTTATTTCACATTACTCACTGTGCGGTATTTCACACCG |
| Mdm10 KO Check | ATGGAGGCCCTTGACATAATCTGAAGTGAA |
| Mdm12 KO F | ACGGTTGAAACAGATCATAAGCTGGCTTCAACTAATCCAAGATTGTACTGAGAGTGCACC |
| Mdm12 KO R | TTTTTTATGTAGACACTATTTTCAAACTATCTTTGTTAAACTGTGCGGTATTTCACACCG |
| Mdm12 KO Check | CGCAATGAGAGGCTTGTTATCCACAGGATT |
| Mdm34 KO F | TAACAAAAAAAACACAACTGCAAGAAACTTCAGCATCGTCGATTGTACTGAGAGTGCACC |
| Mdm34 KO R | ATGTATTTGTGTAGTTATGTACTTAGATATGTAACTTAATCTGTGCGGTATTTCACACCG |
| Mdm34 KO Check | AGGAACAGCTGAACACTATACTGGATACCG |
| Ltc1 KO F | TTG AAC GCA ATG AAG AGA GCA TGG TAC GTT GAA TTG TAG AAG ATT GTA CTG AGA GTG CAC |
| Ltc1 KO R | ACA TAC ATA ACC TTA CAT ACA ATA TTA CAT GAT TAC TCT GCT GTG CGG TAT TTC ACA CCG |

| | |
|---|---|
| Ltc1 KO Check | CGC ACT TGT CAT AGT CGC TGC ACT TGT ATC |
| Vam6 KO F | GCA AAA ACC CTT CAA AAT ATC AAT TTA TAC CAA AAA TTA AAG ATT GTA CTG AGA GTG CAC |
| Vam6 KO R | AAG AAA TAC TAA CAA CAA TAA CAG CAG CTG TTA AGG GAT CCT GTG CGG TAT TTC ACA CCG |
| Vam6 KO Check | CAT TCT GCT GCG CTA CTT CTG TAT TAT TGC |
| Num1 KO F | AAAGACGCAACGGTCAAGGCTTTCCACGAGACGTTCGAATagattgtactgagagtgcac |
| Num1 KO R | TATTGTTCTTAATTTACTTAGAGTTATTTAGTTTTTTTAActgtgcggtatttcacaccg |
| Num1 KO Check | CAAGTGTGGATCCGAGCCCTGTAGTAGAAT |
| Nvj1 KO F | ATCAAAAAAGCTACAAATATAATTGTAAAATATAATAAGCAGATTGTACTGAGAGTGCAC |
| Nvj1 KO R | TAAGTGACGATGATAACCGAGATGACGGAAATATAGTACACTGTGCGGTATTTCACACCG |
| Nvj1 KO Check | TTGGTCACTGCAACTTATACTCCAGGTGAT |
| Gem1 S19N F | GGTTGGTAAAaatAGTCTGATTGTATCATTAAC |
| Gem1 S19N R | CCTTCATCACCGCAAATAAC |
| Gem1 E225K F | AGATGACAACaaaATCTTGGGCTTAC |
| Gem1 E225K R | AAATATGAGTCCTGGTTTAAATC |
| Gem1 E354K F | GAATAATCAAaaaTTACATCGTCTATTTAAG |

| | |
|---|---|
| Gem1 E354K R | AAACCACCATCATTGTCG |
| Gem1 S462N F | TTGCGGCAAAaatTCTTTGCTAGAG |
| Gem S462N R | CATGGCTTTCCAATGACAAAG |
| Chr 1 Integration Check R | GCACTACGTCAATGCAAGT |
| AmpR R | CTG ATC TTC AGC ATC TTT TAC TTT CAC CAG CG |
| Psd1 KO F | GGTCGTTATTTTTTGAAGAAGAAGGAAAAGCAAAGCCAGCAGATT GTACTGAGAGTGCAC |
| Psd1 KO R | TATACAGCAAAATAAATGCTAACTTTACATATGATTGCTTCTGTGC GGTATTTCACACCG |
| Psd1 KO Check | TGACCGTGTTCACTGTGAGCTATTGCAGAA |
| Psd2 KO F | GTA AAG AAT CCT CGA TTT TCA GGA GCA TCC AAC GAC GAA GAG ATT GTA CTG AGA GTG CAC |
| Psd2 KO R | ATT TTG GTA ACC ACT AAC TAC AGC CAA TTT TTC GGC GGC TCT GTG CGG TAT TTC ACA CCG |
| Psd2 KO Check | AAC TAC ACT TGC ATT ATC CTT CCT CGT CCC |
| Ups1 KO F | TGGCTTCTGAGACGGCGGTAAGATATCCTTAAGAGTTGCAagattgta ctgagagtgcac |
| Ups1 KO R | CGCCCATGGTGATATCTTTAAAGATCTTTAAATGGGAACActgtgcggt atttcacaccg |
| Ups1 KO Check | AACCGGAATCAAGCACCAAGGTAGTAAGCA |
| Crd1 KO F | CAGGCCTGGTAGCATAGTTTGGTCCCTAATAATTTAGTCAAGATT GTACTGAGAGTGCAC |

| | |
|---|---|
| Crd1 KO R | TGAAAAGTCAGGACCCTTTTCAAAAAGGATCGCAATTATACTGTG CGGTATTTCACACCG |
| Crd1 KO Check | TATTGGAACTCTGAACCTGTAACAACCAGT |
